# Pro-inflammatory Pathways Contribute to Pathogenesis of *Clostridioides difficile* Infection in a Murine Model - A Spatial Transcriptomics Study

**DOI:** 10.1101/2024.11.15.623793

**Authors:** Niloufar Ghahari, Ahmed AbdelKhalek, Sanjeev Narayanan, Deepti Pillai

**Author notes:** These authors contributed equally.

## Abstract

*Clostridioides difficile* (*C. difficile*) is a common cause of antibiotic-induced diarrhea and causes the highest number of nosocomial infections. Only two antibiotics are currently recommended for treating *C. difficile* infection (CDI), which may contribute to unsatisfactory treatment outcomes and an increased likelihood of recurrence. *Clostridioides difficile* exists as a non-pathogenic member of the human intestinal microbiome in 10-20% of the population, a phenomenon observed in mouse models after infection with bacterial spores. In this study, we aim to evaluate the difference in gene expression between symptomatic and asymptomatic mice after infection with *C. difficile* using spatial transcriptomics analysis. We also aim to evaluate the spatial aspect of altered genes between different layers of intestinal mucosa (superficial vs deep) and identify the key pathways. Formalin-fixed paraffin-embedded (FFPE) intestinal sections were utilized for analysis using NanoString^™^ platform to evaluate differential gene expressions in the caecum and colon. The IL-17 pathway, including *Lcn2, Cxcl2*, and *S100a8* genes, was significantly upregulated in symptomatic mice. The IL-17 signaling pathway activated downstream signaling through NF-κB and MAPK pathways. Gene expression was significantly altered between the intestinal superficial and deep mucosal layers, highlighting layer-specific differences in gene expression patterns in the intestines of symptomatic and asymptomatic mice. Gene expression patterns in the enteric mucosa explain several clinical signs and lesions in CDI mice.

## 1. Introduction

*Clostridioides difficile* is an anaerobic, Gram-positive bacterium that causes severe and life-threatening diarrhea (1). Virulent *C. difficile* strains are highly toxigenic, they secrete two primary enterotoxins; TcdA and TcdB (2,3). These toxins deactivate small GTPases within the host cells after penetrating them, causing disturbances in the actin cytoskeleton, and resulting in the loss of tight junctions in the intestinal epithelial layer (4). This increases permeability and ultimately results in cell death (cytopathic effect) (5). Neutrophil infiltration, a distinctive characteristic of CDI, results in inflammation and mucosal tissue damage (6). CDI occurs when *C. difficile* spores are introduced into the intestines of vulnerable individuals (7). Spores of *C. difficile* can originate either within the body or from external sources, such as contact with contaminated fecal matter (8). In the intestine, the presence of bile salts triggers the germination of *C. difficile* spores into toxin-producing vegetative cells (9). Disturbances in the normal intestinal flora (often due to broad-spectrum antibiotic use, hospitalization, old age, and comorbidities such as HIV or cancer) can lead to *Clostridium difficile* infection (CDI) (10). Although CDI is frequently associated with hospitalization; its pathogenesis remains incompletely understood (11). The Infectious Diseases Society of America (IDSA) and the Society for Healthcare Epidemiology of America (SHEA) limit CDI treatment guidelines to two antibiotics, vancomycin and fidaxomicin. At the same time, metronidazole is only recommended without other agents.(12– 15).

Most host cells possess uniform DNA. Analyzing cells at the protein level rather than the DNA level is more effective in evaluating their variety and functions (16). Current protein profiling technologies offer limited insights into proteins expressed in low abundance and the kinetics of protein expression. Spatial and sequential cross talk between cells at the site of infection can be comprehensively evaluated using mRNA expression signatures (transcriptome) (17). Investigating the contribution of individual cell types could provide novel approaches for preventing and treating *C. difficile* infections. In several previous mouse studies conducted in our and other laboratories, up to 30% of the mice infected with *C. difficile* do not show clinical signs typical to CDI post-infection even though they carry *C. difficile* in their intestinal contents (asymptomatic mice) (18–23). The specific objective of the present study was to determine the differential gene expression in the large intestinal mucosa of the symptomatic mice and compare it to asymptomatic mice using spatial transcriptomics technology. Spatial transcriptomics and direct sequencing of transcripts from the tissue generates a detailed map of diverse cell types and corresponding gene expression patterns in their natural spatial context (24).

As mentioned, enterotoxins are pivotal in the pathogenesis of CDI as they induce a robust inflammatory response in the host (25). Toxin A (TcdA) and toxin B (TcdB) disrupt the cytoskeleton of intestinal epithelial cells by glycosylating Rho family GTPases, leading to cell rounding, loss of barrier function, and cell death (26). This cellular damage triggers the release of pro-inflammatory cytokines and chemokines, which recruit neutrophils and monocytes to the site of infection further perpetuating tissue damage and inflammation. The resulting colonic inflammation manifests as pseudomembranous colitis, characterized by the formation of diphtheritic membrane composed of inflammatory cells, dead epithelial cells, and fibrin (27,28).

## 2. Materials and Methods

### 2.1 Tissue sample collection and processing

Mouse studies were conducted in accordance with the American Veterinary Medical Association (AVMA) guidelines and were approved by the Purdue Institutional Animal Care and Use Com- mittee (IACUC) under protocol number 2008002068. Intestinal tissue samples were from a previous animal study conducted in our laboratory *(23)*]. Briefly, six-week-old mice were adapted to the new environment for one week before oral treatment with a five-antibiotic cocktail for three days. Antibiotic treatment was then ceased for two days to allow for drug clearance before injecting the mice intraperitoneally with clindamycin. One day later, mice were orally infected with *C. difficile* spores. Uninfected mice were used as a control. Mice were monitored for signs of CDI and were euthanized immediately upon the appearance of severe signs. Untreated and asymptomatic mice were euthanized at the end of the experiment using CO_2_ asphyxiation, and organs were collected for histopathological examination and spatial transcriptomic analysis. Thirty colonic and cecal tissues were preserved in 10% neutral buffered formalin, transferred to 70% ethanol and cleared in xylene. Subsequently, the tissues were embedded in paraffin, sectioned, and stained with hematoxylin and eosin (H&E) for histological evaluation. Tissue processing was conducted by the Histology Research Laboratory at Purdue University.

### 2.2 Spatial transcriptomics analysis

NanoString GeoMx spatial profiling platform was utilized. Sections of paraffin embedded FFPE tissues at 5 μm thickness were placed in a positively charged slide (PathSupply, Wilmington, DE, USA). Sections were mounted on microscope slides and dried. The slides were deparaffinized in antigen retrieval solution (Thermo Fisher Scientific) at 100 °C for 20 minutes and treated with 1 µg/mL proteinase K at 37 °C for 15 minutes (Thermo). Mouse transcriptomics was visualized using the GeoMx mouse whole transcriptome atlas (WTA, NanoString, Bruker Spatial Biology, Inc,). An overnight *in situ* hybridization was performed with a mouse WTA probe and the next day the slides were washed twice at 37 °C for 25 minutes with 50% formamide/2X saline-sodium citrate (SSC) buffer to remove unbound probes. The slides were then incubated with morphological markers. Anti-pancytokeratin (Pan-CK) antibody at 1:200 (Thermo) tagged to Alexa Fluor 532, anti-CD45 antibody at 1:200 (CST) tagged to Alexa Fluor 594, and anti-CD3 antibody at 1:150 (OriGene) tagged to AlexaFluor 647 antibodies were used for immunostaining of slides. Nuclei were stained with Syto13 fluorescent dye (Fluorescence 532, Thermo). Stained slides were loaded onto the GeoMx Digital Spatial Profiler (DSP; NanoString) and scanned. After visual inspection using the GeoMx DSP, geometrical regions of interest (ROI) were selected for oligonucleotide collection. Photocleaved oligonucleotides from each spatially resolved ROI were sequenced with the Illumina NextSeq2000 sequencer (Illumina) and downstream analyses were performed using GeoMx Server (NanoString).

### 2.3 Quality control and data normalization

Quality control, normalization, and data analysis were conducted using the GeoMx DSP Analysis Suite (DAS) online portal. After quantifying the probes, the RNA analysis data (DCC files) were transferred to the Nanostring GeoMx platform for detailed QC, normalization, and subsequent analysis. This step facilitated a thorough quality control (QC) assessment using set criteria, the necessary normalization, and analytical procedures. Precise measurement and high-quality data of the RNA samples under study were ensured. As a quality control criterion, the raw read threshold and total number of barcodes were counted by the Illumina sequencer. Only regions with >1000 reads were used for the analysis. Percent aligned reads segments with <80% alignment were flagged for evaluation, assessing potential contamination sources. Sequencing saturation of <50% was flagged. Negative probe count geomean (RNA-NGS) segments of negative probes <10 were flagged, likely to be influenced by slide preparation and wash steps. No template control (NTC) counts of >1000 were analyzed if the DCC file for NTC has low, even count distribution, segments do not have similar deduplicated counts as their NTC, and segments have adequate downstream gene detection. Segments with a cell number or surface area below the threshold were flagged. QC values for % trimmed and % stitched were >90%. Biological QC aimed to remove low-performing probes using one specific probe per gene, focusing on negative probes. A probe was excluded if it was a high or low outlier in >20% of segments. The limit of quantification (LOQ) was defined as two standard deviations above the geometric mean of the negative probes. Segments with low signals were removed retaining segments where 5% of targets were above the expression threshold, set at LOQ below 2. Low-detected genes were removed retaining targets detected in at least 5% of segments as defined by an expression threshold. Technical effects, like segment size and tissue quality, were normalized using third quartile (Q3) normalization. The 75th percentile (Q3) of gene counts was calculated within each segment, and all gene counts were divided by this Q3 value. This step was repeated for each segment, and then all gene counts from all segments were multiplied by the geometric mean of all Q3 counts. Screening genes of interest was performed to eliminate anomalies before consolidating the data into one count per sample and target.

### 2.4 Gene expression analysis

Genes were analyzed across the following comparisons; symptomatic versus asymptomatic mice, superficial layer of symptomatic versus asymptomatic mice, deep layer of symptomatic versus asymptomatic mice, and the superficial versus deep layers of colon, cecum and colon + cecum in both symptomatic and asymptomatic mice. Genes with a Log2 fold change (LFC) of ≥1 were considered significantly upregulated, while genes with an LFC ≤ -1 were considered significantly downregulated. The functions of these significantly dysregulated genes were verified using GeneCards (https://www.genecards.org) and relevant literature.

### 2.5 Pathway analysis and differential expression analysis of data

Pathway and differential expression analyses and differential expression analysis were conducted using the GeoMx Digital Analysis Suite (DAS). The online databases Reactome (https://reactome.org/) and KEGG (https://www.genome.jp/kegg/pathway.html) databases were utilized to analyze and illustrate pathways and pathway maps. Pathways were sorted based on adjusted p-value, coverage, and normalized enrichment score (NES). Pathways with an adjusted p-value of <0.05, coverage equal to or greater than 70%, and a significant NES were prioritized. NES accounts for differences in gene set sizes and variations in enrichment scores across gene sets, providing a normalized measure of pathway enrichment. NES was considered significant if the adjusted p-value was less than 0.05.

### 2.6 Statistical Analysis

Following normalization, we conducted all statistical analyses and created visualizations based on the normalized Log2 counts. To determine expression correlations, we calculated the Pearson correlation coefficient (R) for paired groups, where R >0 indicated a positive correlation and R <0 as a negative correlation. We used linear mixed modeling to assess expression differences between two groups and performed one-way analysis of variance (ANOVA) for comparisons across multiple groups. For the generation of heatmaps, the normalized Log2 counts were zero-centered, and the Euclidean distance method was used to quantify distances. Average hierarchical clustering was then employed to identify expression clusters.

## 3. Results

### 3.1. Selection of ROIs between symptomatic and asymptomatic Mice

Histological differences in the tissue between symptomatic and asymptomatic mice in a CDI mouse model were reported previously (23). Briefly, symptomatic mice exhibited significant edema, neutrophil-driven inflammation, necrosis, and ulceration. The asymptomatic mice had tissue morphology comparable to the uninfected control mice (23). The variation in tissue morphology between symptomatic and asymptomatic mice was analyzed by selecting 25 regions of interest (ROIs) from each sample, using hematoxylin and eosin (H&E), pancytokeratin (PanCK), and CD45 staining. CD3 staining was also performed; however, it did not label sufficient cells to reach the threshold nucleus count, whereas CD45 provided the necessary cellular coverage. ROIs stained positive for pancytokeratin (PanCK+) were classified as areas of illumination (AOIs), indicating epithelial regions. Furthermore, targeted AOIs were analyzed for CD45, a surface marker specific to immune cells such as T cells, B cells, neutrophils, and macrophages (29). Anti-CD45 allowed for the identification and localization of inflammatory cell infiltration within these epithelial regions. CD45 was selected for further analysis to assess immune activity and its correlation with symptomatic versus asymptomatic tissue changes.

Figure 1 shows example ROIs selected from symptomatic and asymptomatic mice samples. These ROIs were chosen from superficial and deep layers of the colon and cecum.

**Figure 1.**
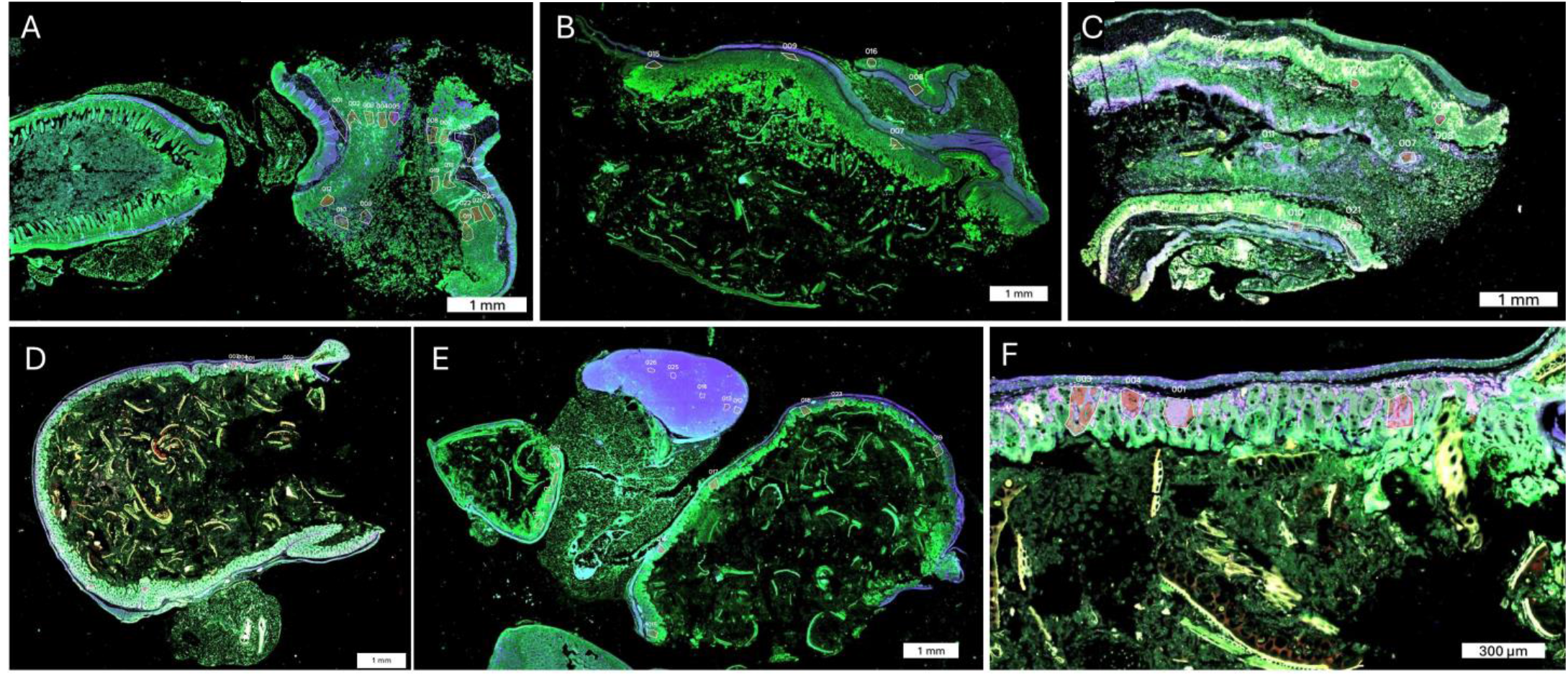
shows example ROIs selected from symptomatic and asymptomatic mice samples. These ROIs were chosen from superficial and deep layers of the colon and cecum. **Figure 1:** A, B, and C depict the regions of interest (ROIs) selected from the superficial and deep layers of the colon and cecum in three symptomatic mice. D and E show the ROIs from the superficial and deep layers of the colon and cecum in two asymptomatic mice. In F, PanCK is shown in red, and CD45 appears in light purple within each selected ROI.

### 3.2 Heat map and cluster analysis result

Figure 2 shows a heatmap displaying cluster analysis of gene expression in *Clostridioides difficil*e-infected mice, comparing symptomatic and asymptomatic groups across the superficial and deep layers of the cecum and colon. The analysis highlights the significant upregulation and downregulation of genes involved in activated inflammatory pathways in symptomatic mice, including Lcn2, Dusp1, Cxcl2, S100a8, S100a9, Defa2, Gstm1, H3c2, H4c17, Cd74, and B2m.

**Figure 2.**
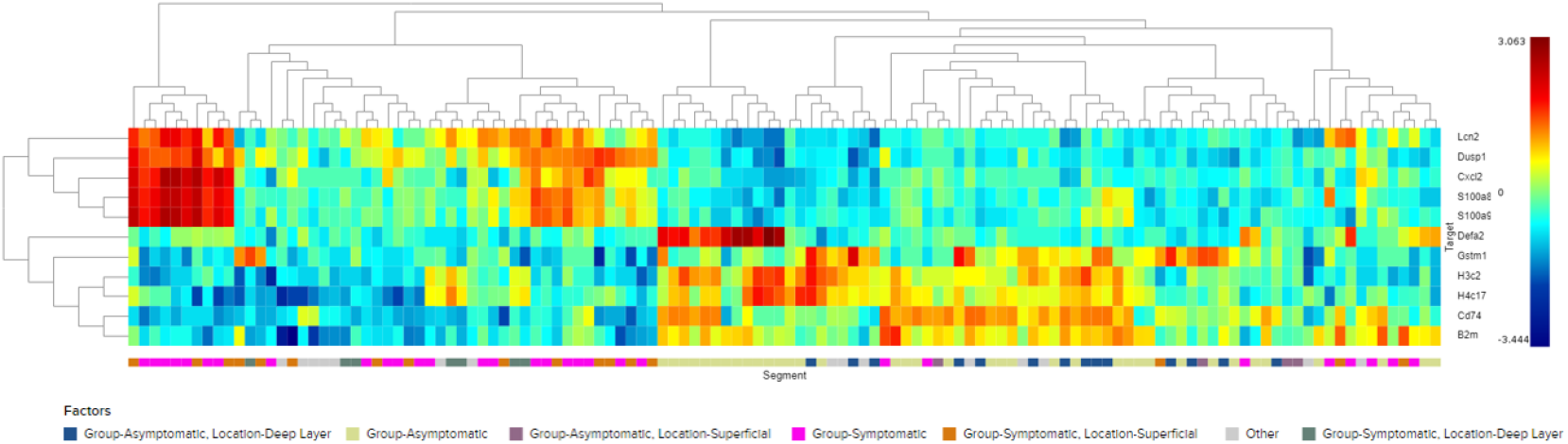
Cluster analysis heatmap displays gene expression in *Clostridioides difficile*-infected mice, comparing symptomatic and asymptomatic groups across superficial and deep layers of the intestine. This visualization distinguishes between significantly upregulated and downregulated genes pivotal to activated pathways in symptomatic mice (*Lcn2, Dusp1, Cxcl2, S100a8, S100a9, Defa2, Gstm1, H3c2, H4c17, Cd74*, and *B2m*). Colors range from red (high expression) to blue (low expression), indicating distinct expression profiles between the groups.

### 3.3 Spatial transcriptomics analysis

#### 3.3.1 Differential transcriptomic analysis of symptomatic versus asymptomatic mice

Significantly upregulated or downregulated genes with known functions were investigated, which was followed by a pathway analysis. The dysregulation of genes was confirmed by performing gene analysis based on the results from pathway analysis. This sequential approach allowed us to transition from pathway analysis to gene-level investigation. Differential gene expression analysis was conducted on RNA samples from selected ROIs within mouse large intestinal mucosa. Upregulated pathways predominantly displayed pro-inflammatory characteristics when comparing all the layers (superficial, middle, and deep) of the colon and cecum of the symptomatic CDI mice to asymptomatic. The interleukin-17 signaling pathway emerged as one of the most upregulated pathways. This pro-inflammatory pathway stimulates other pathways, such as MAPK and NF-κB signaling. Our results indicate a significant upregulation of genes involved in MAPK and NF-κB signaling pathways namely, *CD14, Jun, Areg, Dusp1, Mapk8, Tlr4, Nfkbia*, and *Il1b*.

Figure 3 depicts a volcano plot illustrating the upregulated and downregulated genes in symptomatic CDI mice compared to infected asymptomatic mice. This plot provides a visual representation of the differential gene expression, highlighting the genes that are significantly altered in response to symptomatic infection.

**Figure 3.**
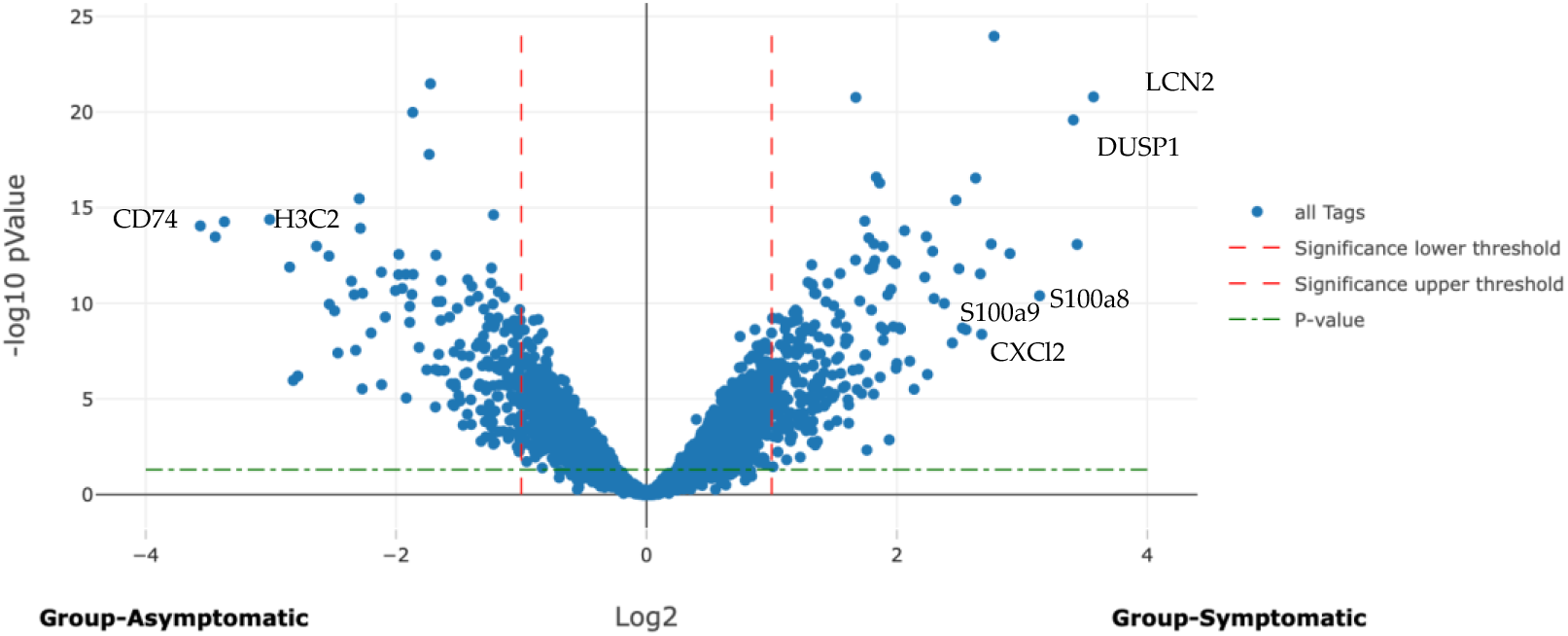
Volcano plot: symptomatic CDI mice upregulated and downregulated genes compared with infected asymptomatic mice, identified 13,801 genes when comparing symptomatic CDI mice to infected asymptomatic mice. Of these, 227 genes had at least a two-fold upregulation in symptomatic mice as evidenced by Log2 fold change (LFC) of ≥1 and 173 genes were significantly downregulated with LFC of ≤-1.

Among the most upregulated genes ere those associated with proinflammatory pathways, particularly the IL-17 signaling pathway. *S100a8* and *S100a9*, crucial genes in IL-17 signaling pathway, were significantly upregulated (LFC= 3.13 and 2.55, respectively). Other upregulated genes include, *Lcn2, Cxcl2*, and *Dusp1* (LFC= 3.56, 2.67 and 3.4, respectively). Genes play a role in gut integrity and mucosal immunity were not significantly upregulated in symptomatic mice, including *Muc1, Ocln*, and *Reg3g*.

Table 2 displays the Log2 fold-change of selected downregulated genes between infected symptomatic and asymptomatic mice. A total of 173 genes were significantly downregulated in symptomatic mice with an LFC of ≤-1, indicating at least a two*-*fold decrease in expression. These values highlight the genes significantly less expressed in symptomatic mice, offering insights into the molecular mechanisms that may contribute to the development of symptoms in CDI. Genes significantly downregulated in comparing symptomatic mice versus asymptomatic mice are mainly associated with crucial cellular processes such as antigen processing and presentation and various metabolic pathways encompassing lipid, amino acid metabolism and PPAR (Peroxisome Proliferator-Activated Receptor) signaling. Within the antigen presentation and processing pathway, several genes including *Cd74, Ctss, Psme2*, and *B2m* showed notable decreases in expression (LFC= -3.56, -1.33, -1.22, and -1.39, respectively). Additionally, significant downregulation was observed in metabolism-related genes, including *Fabp6, Fads2, Hao2*, and *Gstm1* (LFC= -2.46, -1.87, -1.95, and -1.49 respectively). Moreover, genes involved in the neutrophil extracellular trap (NET) formation pathway, including *H3c2* and *H4c17*, also exhibited significant downregulation (LFC= -2.28 and -1.64) in symptomatic mice compared to asymptomatic mice. This suggests a decrease in histone gene expression critical for NET formation. Another significantly downregulated group of genes is the defensin family, including *Defa2, Defa17, Defa34*, and *Defa40* (LFC= -1.46, -1.21, -1.14, and -1.07, respectively), which are part of the IL-17 signaling pathway and contribute to host defense against pathogens.

**Table 1.** Log2 fold change of selected upregulated genes in symptomatic mice. These genes were chosen for their roles in the inflammatory processes and immune response during *Clostridioides difficile* infection (CDI), including those related to the IL-17 signaling pathway and its downstream effects.

| Gene ID | *S100a8* | *S100a9* | *Lcn2* | *Cxcl2* | *Dusp1* |
| --- | --- | --- | --- | --- | --- |
| Log2 fold change | 3.13 | 2.55 | 3.56 | 2.67 | 3.4 |

**Table 2.** Log2 fold-change of selected downregulated genes in infected symptomatic mice. These genes were chosen for their roles in immune defense, antigen processing and presentation, and the inflammatory response during *Clostridioides difficile* infection (CDI), including those related to the IL-17 signaling pathway and its downstream effects.

| Gene ID | *Cd74* | *Gstm1* | *H3c2* | *H4c17* | *B2m* | *Defa2* |
| --- | --- | --- | --- | --- | --- | --- |
| Log2 fold change | -3.56 | -1.49 | -2.28 | -1.39 | -1.64 | -1.46 |

#### 3.3.2 Differential gene expression in superficial and deep intestinal layers between symptomatic and asymptomatic mice

Gene expression was compared in the superficial layer of the colon and cecum of symptomatic mice versus the superficial layers of asymptomatic mice. A similar comparison was made for the deep layer of the colon and cecum. Figure 3 illustrates differential gene expression in the superficial layers and Figure 4 depicts differential gene expression in the deep layers.

**Figure 4.**
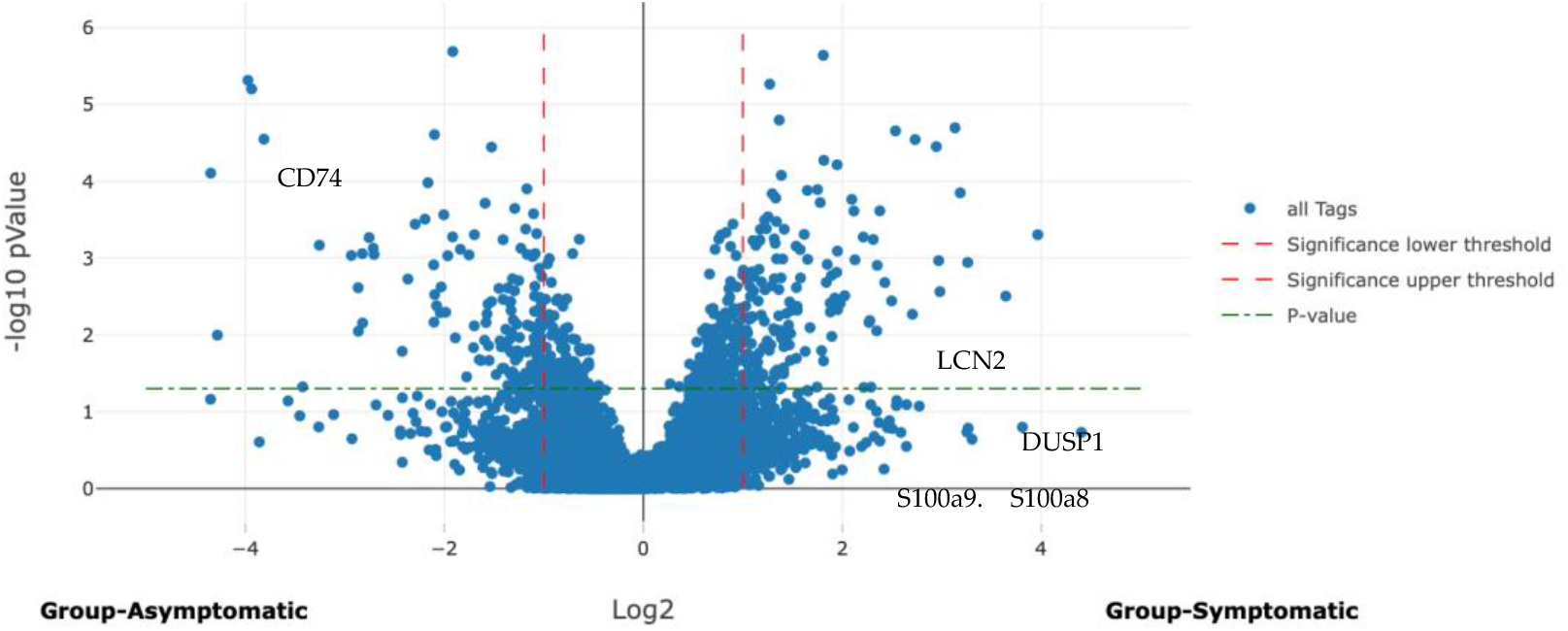
Volcano plot illustrates differential gene expression in the superficial layers of the intestine between symptomatic and asymptomatic mice, identifying the expression of 13,801 genes, 444 genes were significantly upregulated with an LFC of ≥1. Conversely, 506 genes were significantly downregulated with an LFC of ≤-1.

The analysis of differential gene expression between the superficial and deep layers of the colon and cecum in symptomatic mice compared to asymptomatic mice revealed distinct patterns of expression of several genes. The *S100a8* gene was significantly upregulated in both the superficial (LFC= 3.30) and deep layers (LFC= 2.50) of the intestine in symptomatic mice, with higher expression in the superficial layer. *S100a9* also showed significant upregulation in the superficial (LFC= 3.26) and deep layers (LFC= 2.42), with more pronounced expression in the superficial layer. *Dusp1* was significantly upregulated in both layers (superficial LFC= 3.81 and deep LFC= 3.73), showing relatively similar expression. The *Lcn2* gene exhibited significant upregulation in both layers, with higher expression in the deep layer (LFC= 4.44) compared to the superficial layer (LFC= 2.98). *Cxcl2* was significantly upregulated in superficial and deep layers of symptomatic mice (LFC= 2.37 and 1.71, respectively), with greater expression in the superficial layer. The gene expression analysis showed significant downregulation of *Cd74*, Gstm1, B2m, *H3c2*, and *H4c17* in symptomatic mice compared to asymptomatic. Cd74 is notably downregulated in both the superficial and deep layers (LFC= -3.81 and -4.04, respectively). *Gstm1* was also more downregulated in the deep layer (LFC= -3.23) than in the superficial layer (LFC= -1.08). *B2m* showed a similar trend, with more significant downregulation in the superficial layer (LFC= -1.76) than in the deep layer (LFC= -1.13). H3C2 was significantly reduced in the superficial layer (LFC= -1.85), with a minor decrease in the deep layer (LFC= -0.19). *H4c17*, on the other hand, showed significant downregulation in the superficial layer (LFC= -1.53) but a slight increase in the deep layer (LFC= 0.20). These results indicate distinct layer-specific differences in gene expression patterns in the intestines of symptomatic mice and asymptomatic mice.

#### 3.3.3 Differential gene expression between intestinal layers in symptomatic mice

This analysis compared gene expression between the deep and superficial layers in symptomatic mice. Figure 5 shows the differences in gene expression between these layers in symptomatic mice.

**Figure 5.**
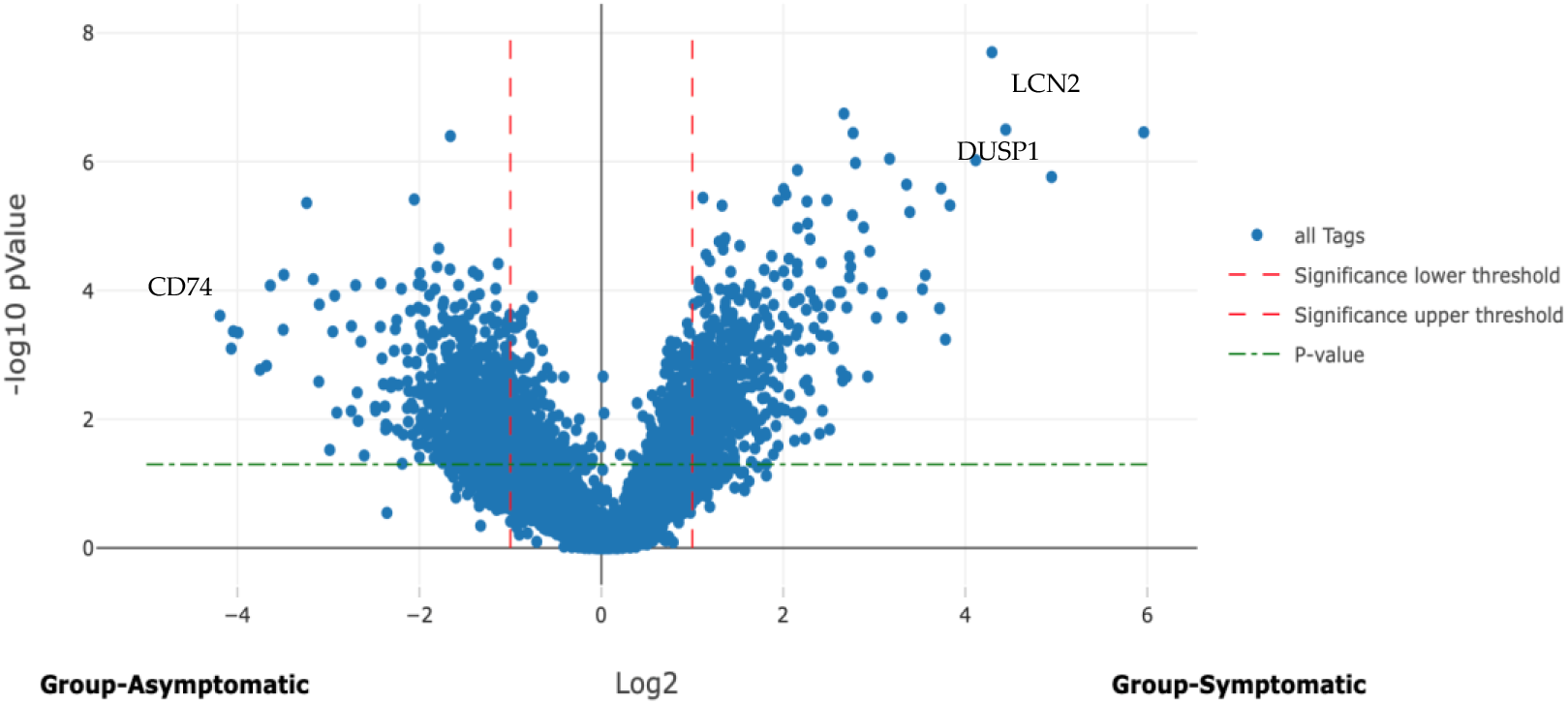
Volcano plot illustrates differential gene expression in the deep layer of the intestine between symptomatic and asymptomatic mice, identifying the expression of 13,801 genes, 663 genes were significantly upregulated with an LFC of ≥1. Conversely, 1833 genes were significantly down-regulated with an LFC of ≤-1.

The analysis of previously identified upregulated and downregulated genes in symptomatic mice compared their expression between the superficial and deep layers of the intestine. For *S100a8*, the LFC was 0.68; for *S100a9*, it was -0.28, indicating no significant changes. *Dusp1* had an LFC of -0.63, while *Lcn2* was -1.32, showing significant downregulation. *Cd74* (LFC= 0.63), *Gstm1* (LFC= 1.52), and *B2m* (LFC= 0.15) varied, with *Gstm1* significantly upregulated. *H3c2* (LFC= -4.00) and *H4c17* (LFC= -3.37) were significantly downregulated. The analysis revealed that genes, including *Gstm1* and *H3c2*, had more pronounced differences in expression profiles.

#### 3.3.4 Differential gene expression between intestinal layers in asymptomatic mice

Figure 6 shows the differences in gene expression between deep and superficial layers in asymptomatic mice.

**Figure 6.**
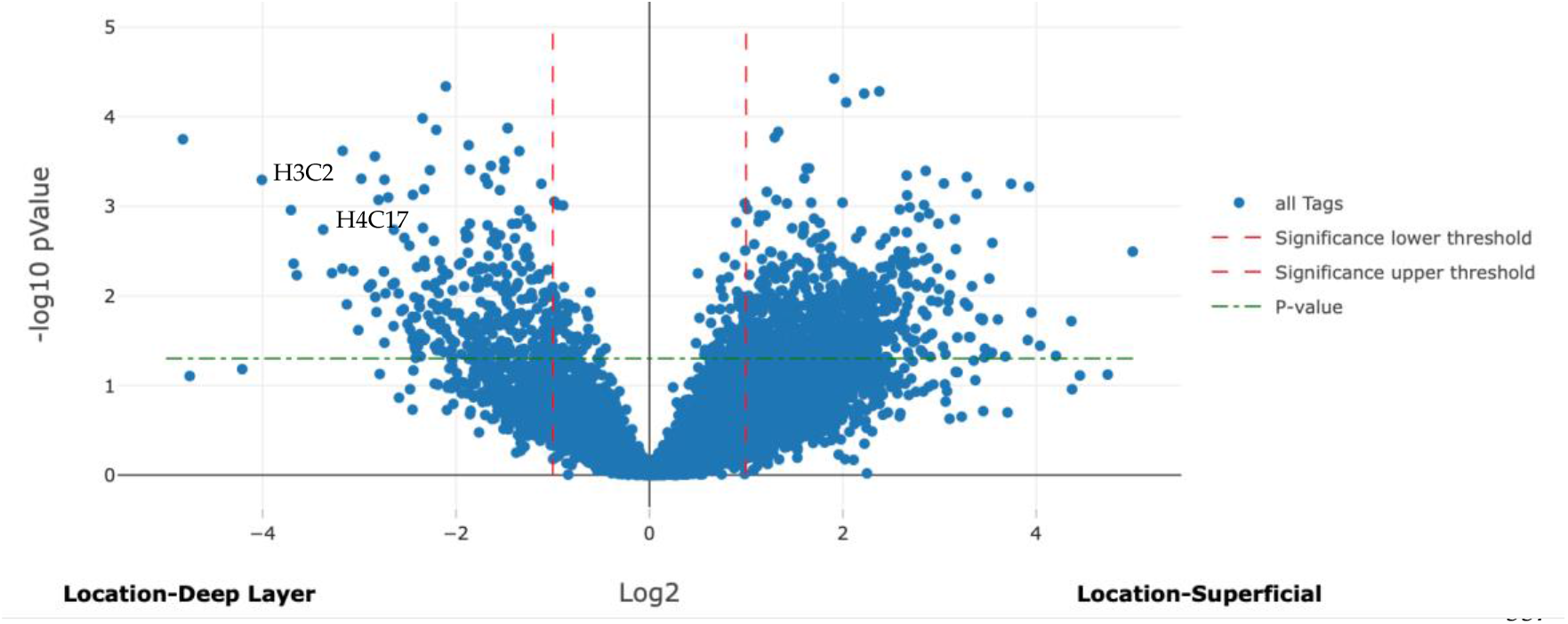
Volcano plot analysis identified 14,025 genes when comparing gene expression between the superficial and deep layers of the intestine in symptomatic mice. Of these, 4,012 genes were upregulated with a Log2 fold change (LFC) of ≥1, indicating at least a two-fold increase in expression in the superficial layer compared to the deep layer. On the other hand, 739 genes were significantly downregulated, with an LFC of ≤-1, reflecting a two-fold decrease in expression in the superficial layer relative to the deep layer. This analysis highlights the differences in gene expression between the two intestinal layers in symptomatic mice.

**Figure 7.**
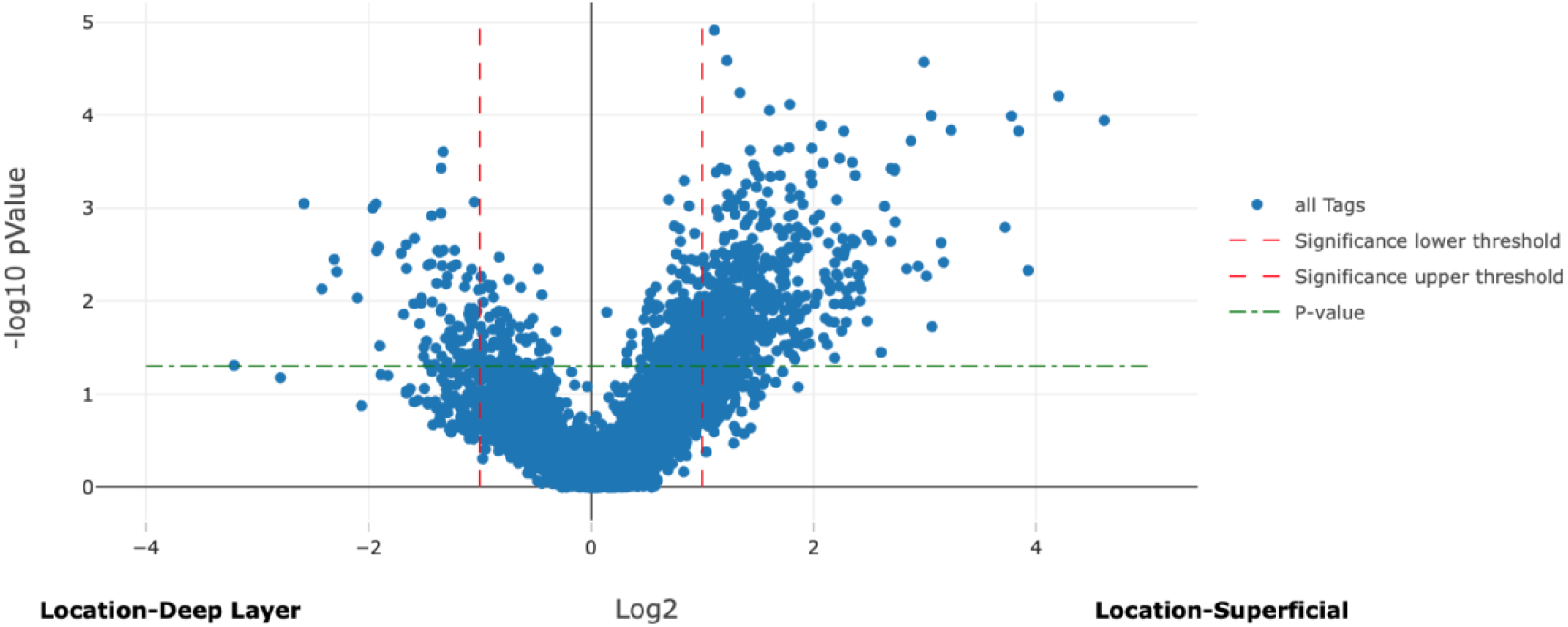
Volcano plot analysis compared gene expression between the superficial and deep layers of the intestine in asymptomatic mice. A total of 12,888 genes were identified in this analysis, 803 genes were significantly upregulated (Log2 fold change [LFC] of ≥1), and 203 genes were significantly downregulated (LFC of ≤-1). This analysis highlighted the differences in gene expression between the two layers of the intestine in asymptomatic mice.

**Figure 8.**
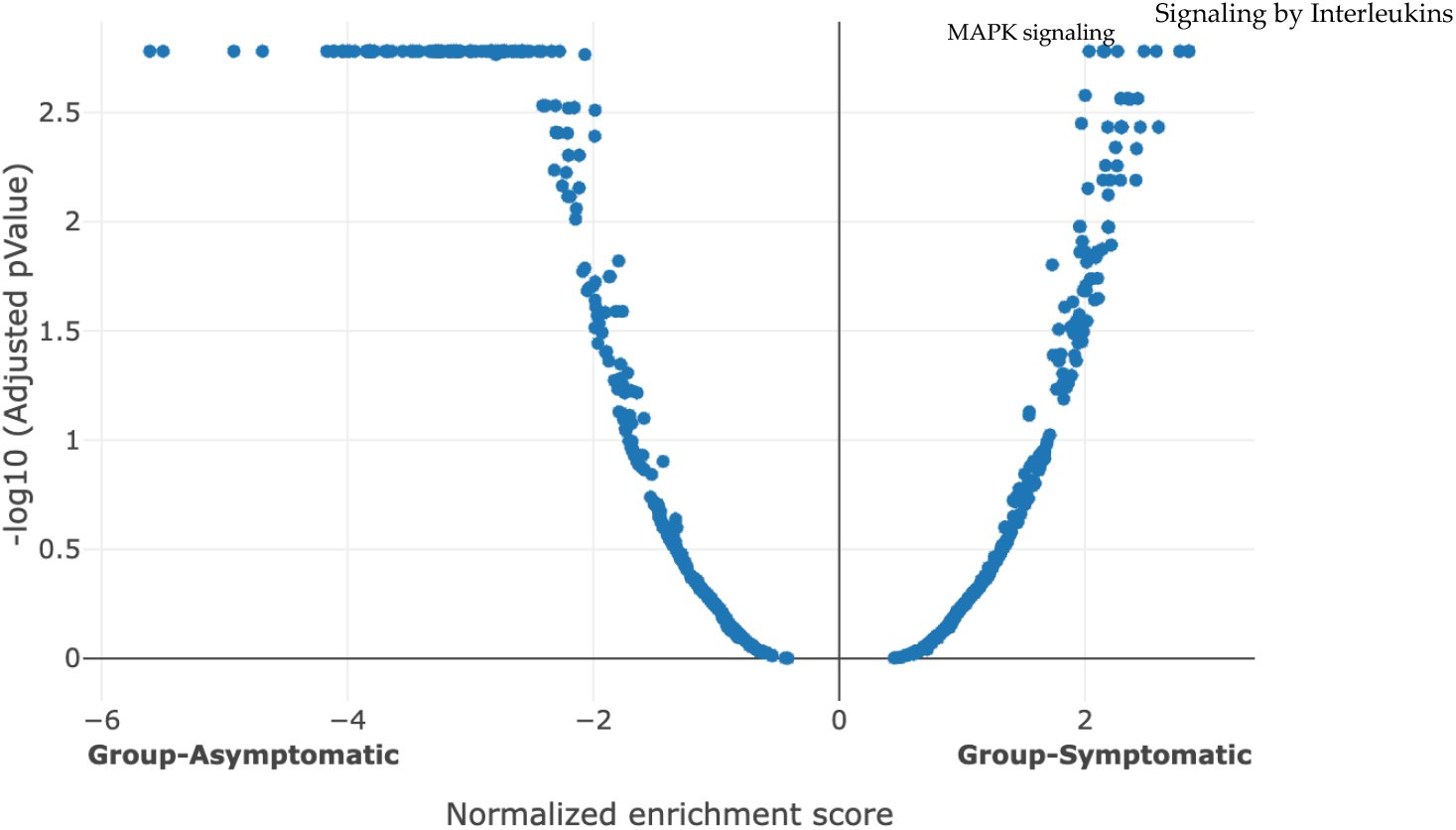
Volcano plot: symptomatic CDI mice upregulated and downregulated pathways compared with infected asymptomatic mice, showing 793 pathways, 350 pathways upregulated, and 443 pathways downregulated.

Gene expression was compared between the superficial and deep layers of the intestine in asymptomatic mice. Out of the genes identified in previous comparisons between symptomatic and asymptomatic conditions, *Cd74, Gstm1, H3c2*, and *H4c17* were significantly downregulated in the superficial layer compared to the deep layer (LFC= -1.10, - 1.50, -1.42 and -1.52, respectively). These differences showcase expression profile nonconformance in gene expression between the intestinal layers in asymptomatic mice.

#### 3.3.5 Pathway Analysis

After individual genes were compared, we were curious to see the difference in pathways between symptomatic and asymptomatic *C. difficile*-infected mice. The volcano plot compared upregulated and downregulated pathways in symptomatic versus asymptomatic mice, mainly highlighting pro-inflammatory pathways. Among the most upregulated pathways were Toll-like receptors, C-type lectin receptors (CLRs), IL-1, and IL-17. We focused on IL-17 because of the significant upregulation of this pathway and genes belonging to this pathway in symptomatic mice. In addition, IL-17 stimulates other pro-inflammatory pathways, including the MAPK signaling pathway and NF-κB signaling pathway. IL-17 signaling pathway was significantly upregulated, covering 100% of the related genes, with a positive normalized enrichment score (NES) and a significant adjusted p-value of 0.018. This pathway activated downstream pathways like MAPK and NF-κB signaling. Some of the upregulated genes in the NF-κB and MAPK signaling pathways include *Cd14, Jun, Mapk8*, and *Areg*, with LFC of 1.38, 1.37, 1.05, and 1.8, respectively. The MAPK signaling pathway had 80% coverage, a positive NES, and an adjusted p-value of 0.01. The TRAF-mediated NF-κB and MAP kinase activation pathways showed 98.55% coverage, positive NES, and a significant adjusted p-value of 0.003. Conversely, pathways significantly downregulated in symptomatic mice included antigen processing and presentation, metabolic pathways for lipids, amino acids, and the PPAR signaling pathway. These pathways had negative NES values and p-values below 0.05. Specifically, the MHC class II antigen presentation pathway had a significant downregulation, with a p-value of 0.001. Similarly, the antigen processing-cross presentation pathway showed significant downregulation with a negative NES and a p-value of 0.03.

## 4. Discussion

The primary goal of this research was to explore the functional and spatial roles of genes and pathways altered in symptomatic mice to understand *C. difficile* pathogenesis, particularly the inflammatory pathways’ contribution to the clinical signs and lesions of CDI. This study investigated the differential gene expression in the intestinal mucosa of mice infected with *C. difficile*. Symptomatic infected mice exhibited altered gene expression compared to the asymptomatic mice. Generally, inflammatory mediators related to epithelial damage and ulceration were found to be upregulated in the symptomatic mice (e.g. *Cxcl2, Lcn2*, and NF-kB). In addition, pro-inflammatory pathways (e.g. IL-17) were significantly upregulated in symptomatic mice (30–32).

The IL-17 pathway is significantly upregulated in symptomatic mice, playing a critical role in host defense against infections (33). This pathway is essential for recruiting immune cells, like neutrophils, to the site of infection and regulating inflammation (34–37). The role of IL-17 in *Clostridioides difficile* infection (CDI) has been studied, illustrating its involvement in both host defense and exacerbation of disease. A study on cytokine profiles in CDI patients found that IL-17 levels were significantly elevated in severe cases, with a shift from Th1 to Th17 dominance observed in these patients. This shift suggests that IL-17 and the Th17 response contribute to the intense inflammation and tissue damage characteristic of advanced CDI, marking IL-17 as both a severity marker and a potential cause of pathogenic immune responses in severe cases (38). Another study expanded on these findings by examining IL-17 production by γδ T cells, showing that these cells are a primary source of IL-17 during CDI and play a protective role in the immune response. In murine models, IL-17-producing γδ T cells were essential for recruiting neutrophils to infection sites, facilitating pathogen clearance while managing inflammation. Mice deficient in γδ T cells exhibited increased susceptibility to CDI, indicating that IL-17 from γδ T cells is crucial for effective host defense, especially in early infection stages (36). A recent study focused on IL-17’s role in recurrent CDI (RCDI), where elevated IL-17 levels were observed in an RCDI mouse model. Treatment with a RORγt inhibitor to block Th17 function significantly improved clinical signs, including reduced inflammation and restored gut barrier integrity. These findings show IL-17’s role in perpetuating inflammation in recurrent cases and suggest that targeting Th17 pathways may offer therapeutic potential for managing RCDI (37). In this study, chemokines like *Cxcl2* were upregulated, reflecting active recruitment of immune cells in symptomatic mice. Molecules such as defensins, occluding (*Ocln*), and *Reg3g*, which are involved in the IL-17 signaling pathway, showed significant differences in expression in expression in symptomatic mice versus asymptomatic mice. Defensins were notably downregulated, indicating a weakened innate immune defense, while *Reg3g* and *Ocln* were upregulated, reinforcing gut barrier integrity (39). *Muc1*, typically involved in mucosal barrier protection (40,41), did not show significant changes in this study, suggesting its role may not be as prominent. The study showed that while some mucosal immunity genes, like *Reg3g* and *Ocln*, were more highly expressed in the superficial intestinal layers of symptomatic mice, overall changes across all layers were insignificant. The significant downregulation of defensins in symptomatic mice likely weakens the host’s ability to fight bacterial infections, particularly in CDI (42). The study showed that the general decrease in defensin expression compromises the host’s defense mechanisms despite localized upregulation of some immune-related genes (43). Since IL-17 is a critical pathway in inflammatory bowel disease (IBD) and ulcerative colitis (UC) (44), it is currently targeted via monoclonal antibodies (e.g. secukinomab) for these conditions (45,46). These findings suggest that targeting the IL-17 signaling pathway, particularly at the level of receptors or key signaling molecules like *Traf6*, could offer an effective therapeutic strategy for CDI. It remains unclear whether the upregulation of the IL-17 pathway observed in symptomatic mice directly results from increased neutrophil infiltration or if the pathway itself induces neutrophil recruitment. This distinction is critical for understanding the exact role of IL-17 signaling in CDI pathology and could influence future therapeutic strategies to modulate the inflammatory response.

The upregulation of *Cxcl2* in symptomatic mice likely contributes to the excessive neutrophil infiltration seen in the inflamed intestinal tissue, exacerbating mucosal damage and contributing to the pathology of the disease (47). While neutrophil recruitment is essential for pathogen clearance, it can also lead to tissue injury, as seen in the characteristic signs of pseudomembranous colitis (48). Blocking *Cxcl2* or its associated pathways could be a potential therapeutic strategy to reduce neutrophil-driven damage while preserving the immune system’s ability to combat *C. difficile* infection, potentially improving disease outcomes in CDI. Lipocalin-2 (*Lcn2*) was also upregulated in symptomatic mice. *Lcn2* is a glycoprotein involved in the innate immune response and inflammation, particularly through the IL-17 signaling pathway (49). The role of *Lcn2* in neutrophil recruitment and inflammation (50) suggests its relevance to the clinical signs of CDI, where neutrophil-driven tissue damage is a hallmark. The increased expression of *Lcn2* in symptomatic mice indicates its contribution to the inflammatory processes that exacerbate CDI. Given that *Lcn2* is also involved in inflammation and cancer progression (51–53), it is plausible that its upregulation enhances neutrophil activity and perpetuates the inflammatory response seen in CDI.

*S100a8* and *S100a9* were among the most upregulated genes in symptomatic mice, correlating with the clinical signs of CDI. The upregulation of *S100a8* and *S100a9* supports the inflammatory response in CDI, as these genes are known to play a crucial role in neutrophil recruitment and activation (54). Their increased expression in symptomatic mice suggests that they significantly contribute to the neutrophil-driven immune response, leading to tissue damage and exacerbating the clinical manifestations of CDI. The increased activity of these genes amplifies the inflammatory lesions characteristic of the infection, highlighting their roles in the disease’s progression and severity. The S100 protein family, particularly *S100a8*, is essential for regulating cell cycle progression and differentiation. This protein acts as a danger-associated molecular pattern (DAMP), crucial in alerting the innate immune system (55). It triggers immune responses by interacting with pattern recognition receptors such as Toll-like receptor 4 (TLR4) and the receptor for advanced glycation end products (*Ager*) (56). *Ager* was significantly upregulated when comparing symptomatic and asymptomatic mice. Micro- or macro-molecules that block the function of these molecules might provide an effective approach to mitigate CDI symptom onset.

Some genes with an anti-inflammatory function were also upregulated including *Dusp1. Dusp1*, known as dual-specificity phosphatase-1 or MAPK phosphatase-1, and is a negative regulator of MAPK activity (57). Upregulation of Dusp1 was also seen in our previous study (23). It is part of the MAPK phosphatase family and is crucial for antiinflammatory responses (58). The increased expression of *Dusp1* in symptomatic mice indicates an attempt by the host to activate anti-inflammatory responses. However, this response seems inadequate as the overall MAPK signaling pathway activity, which typically drives inflammation, remains elevated in symptomatic mice compared to asymptomatic ones. Research involving the silencing of the *Dusp1* gene using lentiviral vector-mediated siRNA in mice with acute pancreatitis demonstrated a higher release of proinflammatory cytokines (59).

The expression of genes involved in the arachidonic acid pathway did not change significantly in this study. A human study found that patients who received NSAIDs had slightly lower mortality rates than those who did not; however, this difference was not statistically significant (60). We aimed to explore the effects of NSAIDs in a mouse model. In the current study, *Fabp6* and *Fads2* were significantly downregulated in symptomatic mice, while *Fabp4* was significantly upregulated. These genes, involved in lipid metabolism and transport, are linked to the arachidonic acid pathway by regulating the availability of fatty acids for inflammatory mediators (61). The downregulation of *Fabp6* and *Fads2* suggests a disruption in lipid metabolism, potentially exacerbating the inflammatory environment, while the upregulation of *Fabp4* indicates a possible compensatory mechanism in lipid handling in symptomatic mice. The downregulation of these genes suggests a disruption in lipid metabolism, potentially exacerbating the inflammatory environment in symptomatic mice, which may have contributed to the worsened outcomes observed in the prior NSAID treatment study.

*Cd74* expression was significantly downregulated in symptomatic mice compared to asymptomatic mice, which may have important implications for the immune response during CDI. *Cd74* is essential for proper immune function, particularly in antigen-presenting cells (APCs) (62), where it acts as a chaperone for MHC class II molecules and supports the activation of immune pathways like NF-κB and ERK (63). This reduction in *Cd74* may impair immune surveillance and hinder the host’s ability to present antigens to clear the infection effectively. The decreased expression of *Cd74* in symptomatic mice suggests that the immune response is compromised, potentially contributing to the severity of CDI. While reduced *Cd74* might prevent excessive inflammation in some contexts, it may also weaken the ability to eliminate pathogens during infection (64).

*B2m* gene was downregulated in symptomatic mice, which likely plays a role in the impaired immune response observed during CDI. *B2m* encodes beta-2 microglobulin, an essential component of MHC class I molecules for presenting antigens to cytotoxic T cells (65). This process is vital for immune surveillance and for efficiently eliminating infected cells (66). The downregulation of *B2m* may reduce the ability to present antigens, weakening the host’s immune defense against infection. This immune dysfunction could exacerbate the severity of the disease. The downregulation of *Cd74* and *B2m* suggests a broad impairment in the immune system’s ability to present antigens and mount an effective immune response. Enhancing *Cd74* and *B2m* expression could be a beneficial therapeutic strategy, improving antigen presentation and pathogen clearance in CDI, thereby mitigating the disease severity.

*Gstm1* gene was downregulated in symptomatic mice, likely contributing to the increased oxidative stress observed during CDI. *Gstm1* encodes the enzyme glutathione S-transferase M1, a key player in detoxification of free radicals and regulating oxidative stress (67). This enzyme neutralizes reactive oxygen species (ROS) and harmful electrophilic compounds that can damage cells and tissues (68). Oxidative stress and free radical damage are significant contributors to the pathophysiology of CDI (69) and the decreased expression of *Gstm1* partially explains the shift toward a pro-oxidative state in symptomatic mice. This shift likely worsens the inflammatory damage to the mucosa, leading to the clinical signs observed in CDI, such as tissue necrosis and ulceration. Restoring *Gstm1* expression or targeting oxidative stress pathways could offer therapeutic potential to mitigate the damage caused by ROS and improve recovery in CDI.

*H3c2* and *H4c17* genes were downregulated in symptomatic mice, impacting the formation of neutrophil extracellular traps (NETs) during CDI. NETs, composed of decondensed chromatin and histones such as H3 and H4, play a crucial role in the immune response by trapping (70) and neutralizing pathogens like *C. difficile*. The downregulation of these histone genes in symptomatic mice could compromise the ability of acute inflammatory response to sequester the infectious agent, allowing *C. difficile* to evade immune responses and exacerbate inflammation. This reduction in NET formation might explain the heightened infection severity and tissue damage observed in symptomatic mice (6). Targeting pathways involved in NET formation or enhancing the expression of H3C2 and H4C17 may represent potential therapeutic strategies to boost immune defenses and improve pathogen clearance in CDI. Based on the findings of this study, significant differences in gene expression were observed between symptomatic and asymptomatic mice. These differences could explain why some mice remained asymptomatic. Several factors might have influenced this, including variability in host responses and immune regulatory mechanisms. Differences in immune responses to the pathogens along with variations in gene expression and pathway activation, could have contributed to the differences in disease presentation, even under identical experimental conditions. Another important factor that could have played a role is the gut microbiome. However, this study did not include the gut microbiome factor as a variable. Given the identical conditions of these inbred mice, this would be surprising.

One of the study’s limitations is the lack of RT-PCR confirmation for the up-regulation and down-regulation of the selected genes. Future research on managing CDI will incorporate RT-PCR as a control measure to validate these findings.

## 5. Conclusions

Our research elucidates the intricate interactions of various genes and pathways contributing to the pathology attribute to *Clostridioides difficile* infection (CDI). We identified several genes and pathways linked to inflammation, immune response, and cellular metabolism by comparing gene expression profiles between symptomatic and asymptomatic mice post-CDI. Disruption in these genes impair the host’s capacity to mount effective inflammatory and immune responses, leading to heightened susceptibility to CDI-induced tissue damage and exacerbated clinical signs. This study is the first to evaluate differential gene expression between the superficial and deep layers of the intestinal mucosa. Our findings reveal specific genes are altered in the superficial versus deep intestinal mucosal layers after CDI. This dimensional analysis provides deeper insights into the spatial regulation of gene expression in the context of CDI. Future research will focus on targeted interventions to modulate expression of these genes and pathways or their activity, which could pave the way for innovative therapeutic strategies against CDI. Given that some upregulated pathways, such as the IL-17 signaling pathway, have pro-inflammatory and anti-inflammatory roles, our approach will focus on precisely targeting select upregulated genes rather than broadly suppressing entire pathways. This refined strategy aims to effectively reduce CDI symptoms while preserving the beneficial aspects of these pathways, offering more targeted therapeutics.

